# Microbial Cell-Adaptive Segmentation in Quantitative Phase Imaging

**DOI:** 10.1101/2025.11.13.688197

**Authors:** Sabrina J. Tuson, Shahla Nemati, Alexander B. Alleman, Christopher J. Marx, Andreas E. Vasdekis

**Affiliations:** Department of Physics, University of Idaho, 875 Perimeter Drive, Moscow ID 83844, USA; Department of Biological Sciences, University of Idaho, 875 Perimeter Drive, Moscow ID 83844, USA; Department of Natural and Environmental Sciences, Western Colorado University, 1 Western Way, Gunnison CO 81231, USA

## Abstract

Quantitative phase imaging (QPI) enables label-free microscopy with exceptional signal-to-background contrast and provides direct measurements of key biophysical parameters such as cell mass. Despite these advantages, QPI lacks a generalized, accurate, and computationally efficient segmentation method, limiting its use in high-throughput single-cell biology. Here, we introduce microbial Cell-Adaptive Segmentation (mCAS), a purpose-built framework for QPI. mCAS is an object-based, modular pipeline that leverages the robust phase-edge signal to achieve accurate segmentation without Fourier filtering, deconvolution, or deep learning. It runs efficiently on standard hardware, is readily implemented in common software platforms, and requires no curated training data. mCAS consistently outperformed both intensity-based and deep learning approaches, achieving over 98% accuracy with up to tenfold faster processing. By eliminating the segmentation bottleneck, mCAS enables scalable, high-fidelity analysis of QPI datasets and provides a broadly applicable foundation for label-free, high-throughput single-cell biology. We validate its generality across diverse bacterial species, morphologies, and intracellular complexities.

**Significance Statement:** Quantitative phase imaging (QPI) enables label-free, noninvasive measurement of cellular biophysics, yet its broader adoption has been limited by the absence of a generalized and computationally efficient segmentation strategy. We present microbial Cell-Adaptive Segmentation (mCAS), a framework purpose-built for QPI that achieves high accuracy without training, specialized hardware, or intensive preprocessing. mCAS generalizes across microbial species and morphologies, outperforming state-of-the-art deep learning approaches while processing thousands of single-cell images in minutes. By eliminating the segmentation bottleneck, mCAS makes QPI broadly accessible for high-throughput, label-free single-cell biology and expands the computational toolkit for quantitative imaging in microbiology.

## Introduction

High-throughput microscopy can capture thousands of high-resolution images in minutes, but converting these data into meaningful quantitative single-cell biology results remains limited by image-processing bottlenecks. Key among these is cell segmentation, which defines cell boundaries and accelerates downstream analyses. [1] Robust segmentation methods must minimize error, scale to large datasets, and adapt across cell types and imaging conditions. A variety of pipelines have been developed for fluorescence and phase-contrast imaging of microbial cells, such as *CellProfiler*, [2–3] *Oufti*, [4] *FogBank*, [5] and *CellTracer*, [6] as well as active contouring methods. [7] More recently, deep learning (DL) approaches have demonstrated impressive versatility, [8–10] yet they require curated training datasets, retraining to handle new conditions, and elevated computational costs.

In contrast, quantitative phase imaging (QPI) lacks generalized segmentation tools, despite its rapidly growing role in single-cell biology [11–17]. This limit is particularly pertinent to microbial cells, which, unlike their mammalian counterparts, [18–19] are considerably smaller, approximately at the resolution limit of magnification objectives such as 40×, making segmentation particularly challenging. Generally, QPI is an interferometric, label-free modality that produces high signal-to-background images by interfering a sample-transmitted beam with a reference beam (**Fig. 1**). [20] Unlike fluorescence, QPI provides direct measurements of dry-mass at low irradiance levels, making it ideal for long-term live-cell studies and high-throughput phenotyping. Yet, the very strengths of optical transparency and label-free imaging, also pose unique segmentation challenges. Halos, background variation, and heterogeneous intracellular contrast frequently limit conventional intensity-based methods. Previous strategies have been restricted to narrow-use-cases, such as segmenting small colonies [21] or large colonies with halo correction and deconvolution [16, 22–23] that remain computationally demanding and often require manual curation. To date, no generalized segmentation method exists for microbial cells in QPI that can operate reliably across diverse morphologies and intracellular complexities without heavy computational cost. [14,18–19,21]

**Figure 1:**
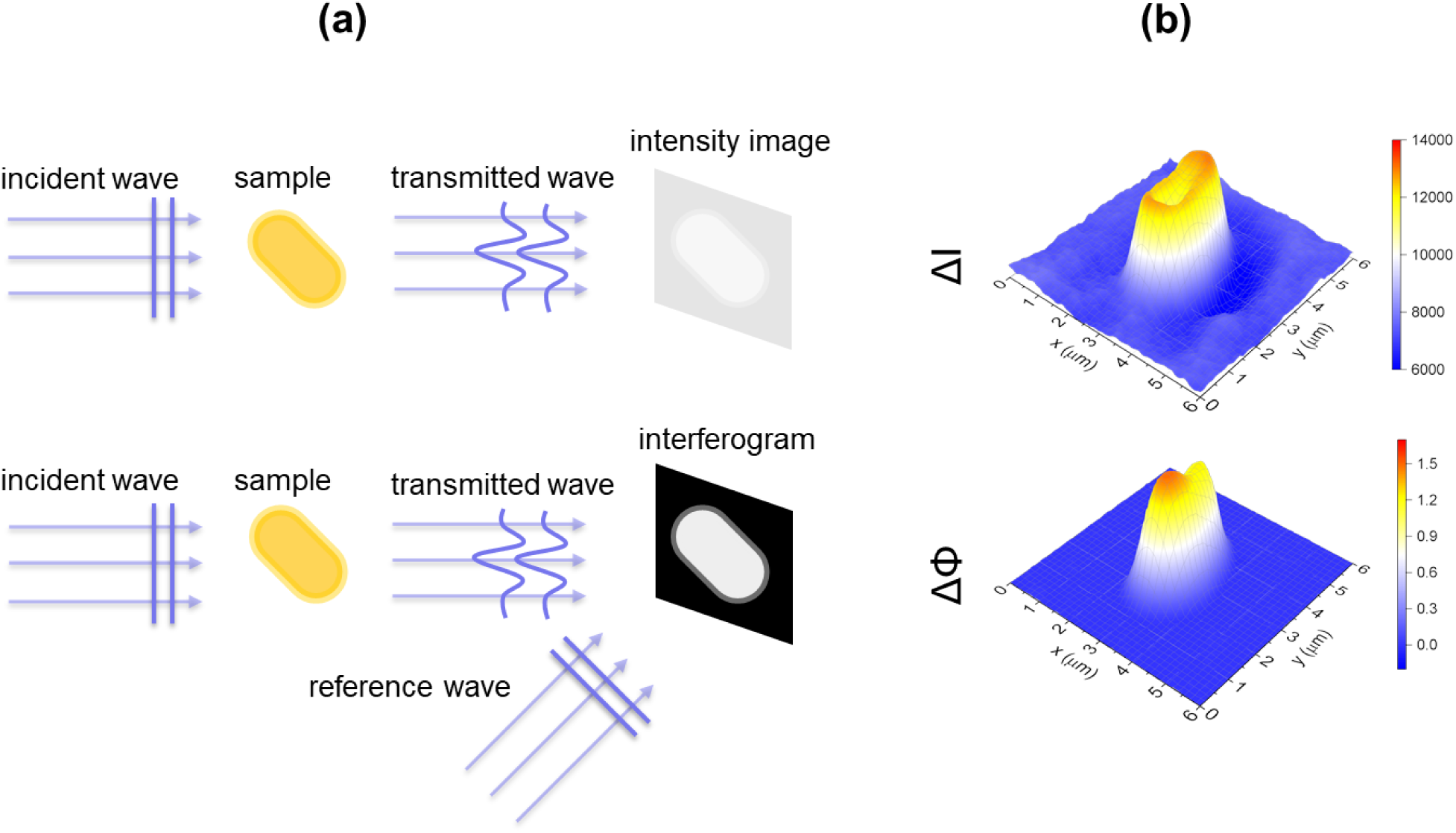
QPI provides enhanced contrast over intensity-based microscopy. **(a)** QPI interferograms are generated by combining the transmitted wave from a sample with a reference wave that has not interacted with the sample, enabling high-contrast visualization of transparent objects. **(b)** A single *Methylobacterium extorquens* cell imaged by phase contrast (*top*) and QPI (*bottom*). In phase contrast, image contrast arises from intensity differences (ΔI), whereas in QPI it arises from phase-delay differences (ΔΦ), producing a sharper signal and minimal background.

To address this technological gap, we introduce *microbial Cell-Adaptive Segmentation* (mCAS), a generalized segmentation framework purpose-built for QPI. mCAS is an object-driven, modular pipeline that exploits the robust phase-edge signal to achieve accurate boundary detection without Fourier filtering, deconvolution, or DL training. Its customizable morphological and contour-shaping steps allow rapid adaptation to different species, morphologies, and imaging conditions. mCAS runs efficiently on standard desktop hardware, is implemented in widely used platforms, such as ImageJ, Python, and MATLAB, and requires no curated training datasets. Through direct benchmarking, we demonstrate that mCAS outperforms both traditional thresholding and state-of-the-art DL-based methods, establishing a versatile, high-performance solution for label-free microbial cell segmentation in QPI.

## Results

### Segmentation Procedure

Unlike conventional segmentation methods that rely primarily on intensity thresholding to classify pixels as object or background, mCAS is driven by edge detection. [24–25] Edge detection identifies object boundaries via abrupt pixel-intensity changes, classifying pixels as edge or non-edge. Because edge-detection is widely used in computer vision and DL, many robust implementations already exist. Accordingly, we implemented mCAS in multiple programming environments, including ImageJ and FIJI, as well as Python and MATLAB, ensuring reproducibility and broad accessibility. [26–27] The mCAS workflow consists of six sequential steps (**Fig. 2a**):

1. **Sobel edge detection** of raw QPI data. [24]
2. **Otsu’s thresholding** to binarize the edge image. [28]
3. **Binary hole filling** to restore enclosed interiors.
4. **Adjustable binary erosion** to remove residual noise.
5. **Adjustable watershed segmentation** at σ = 1.5. [29]
6. **Cell-mask shaping** using bounding box or spline fitting.

**Figure 2:**
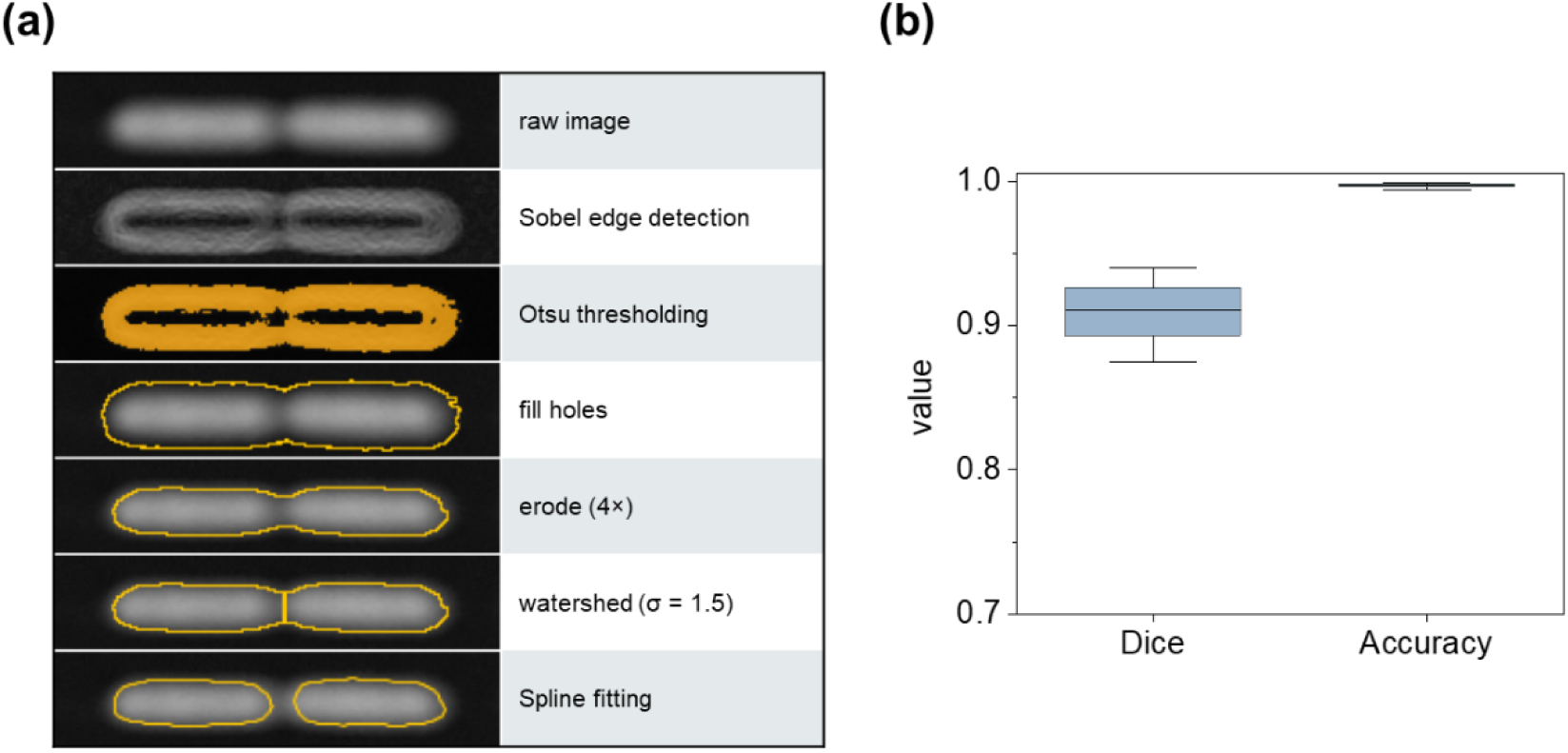
mCAS workflow and metrics. **(a)** The mCAS workflow consists of the following steps (1) Sobel edge detection to enhance phase boundaries. (2) Otsu’s thresholding to convert the edge image to a binary mask. (3) Binary hole filling to restore enclosed interiors. (4) Adjustable binary erosion to remove residual noise. (5) Adjustable watershed to separate touching cells using a tunable geometric criterion. (6) Cell-mask shaping to produce the final segmentation output. **(b)** Dice accuracy metrics for mCAS-generated masks compared with ground truth fluorescent images of *E. coli* expressing tdCherry fluorescent protein. [31] The difference between the Dice and accuracy metrics can be attributed to a higher contribution of background (true negatives) to the accuracy score (n = 350); box charts represent 25-75% of the individual scores; whiskers indicate 10-90% of the individual scores; and horizontal lines indicate the median score.

We found that this segmentation pipeline produced high-fidelity masks, with an average accuracy metric exceeding 0.99 when compared to masks generated by thresholding standard fluorescence intensity images of the same cells expressing fluorescent proteins (**Fig. 2b**). This performance improvement stems in part from the Sobel operator, which computes discrete image gradients in both horizontal and vertical directions. In evaluating other common edge detectors (**Fig. S1**), we found Sobel and Prewitt to be preferable for balancing high fidelity boundary localization while maintaining computational efficiency, due to their use of only two kernels. [24] We therefore adopted Sobel [24] for subsequent analyses, while noting that Prewitt offered comparable performance.

To convert the edge-enhanced image into a binary mask, we combined Otsu’s thresholding [28] with binary hole filling. Otsu’s threshold minimized segmentation errors relative to other approaches, though this step is modular and alternatives, such as maximum entropy thresholding, can be substituted easily. Since we performed thresholding on an edge-enhanced image, object interiors often retained low gradient values and may be misclassified as background. We corrected this by applying binary hole filling. This step reclassified fully enclosed, four-connected background pixels as foreground, [30] yielding contiguous segmented regions suitable for downstream analysis. Following hole filling, we applied binary erosion to refine boundaries by removing residual noise and disconnected artifacts. The extent of erosion is adjustable and depends on image quality, object morphology, and the algorithm used by the platform. Further, making this step adjustable enhances mCAS’s adaptability to diverse datasets while preserving biologically relevant features such as cell size and circularity.

To resolve touching or overlapping cells, mCAS employs common watershed strategies. We used ImageJ’s distance-transform, which converts the binary mask into a grayscale distance map. In this context, pixel values corresponded to the distance from background pixels. Local maxima served as watershed seeds, and regions expanded until they collide with object edges or neighboring regions. When collisions occur, objects are separated according to a tunable geometric criterion: the ratio between the neck radius (the narrowest point between two objects) and the radius of the larger object. [30] The adjustable watershed plugin [29] enabled us to tune this splitting threshold, ensuring accurate segmentation in densely packed or irregularly shaped populations, critical for high-throughput imaging workflows.

As a final step, mCAS incorporates cell-mask shaping. Here, we evaluated two approaches: bounding box fitting and polynomial spline fitting. In bounding box shaping, the bounds of the object are first determined and then the shortest ends of the bounding box are replaced with half-circles with radii equal to their original side length, ultimately producing a model rod shape. This shaping method is computationally efficient and well-suited for simple rod-shaped morphologies such as *Escherichia coli*. However, it lacks flexibility for curved or irregular shapes. In contrast, spline fitting provided more accurate outlines for complex morphologies, such as the irregular contours of *Methylobacterium extorquens* imaged on agarose pads. This method involved fitting polynomial splines to contours generated by watershed segmentation. A critical parameter is the sampling rate of contour points: oversampling produced pixelated outlines, while undersampling yielded overly smooth contours that deviate from true boundaries. To address this, we developed a Python script that empirically optimized sampling rates, balancing computational efficiency with biological accuracy. This ensured high-quality reconstructions of cell shapes that enabled robust quantification of descriptors such as perimeter, elongation, and curvature (**Fig. 3a**).

**Figure 3:**
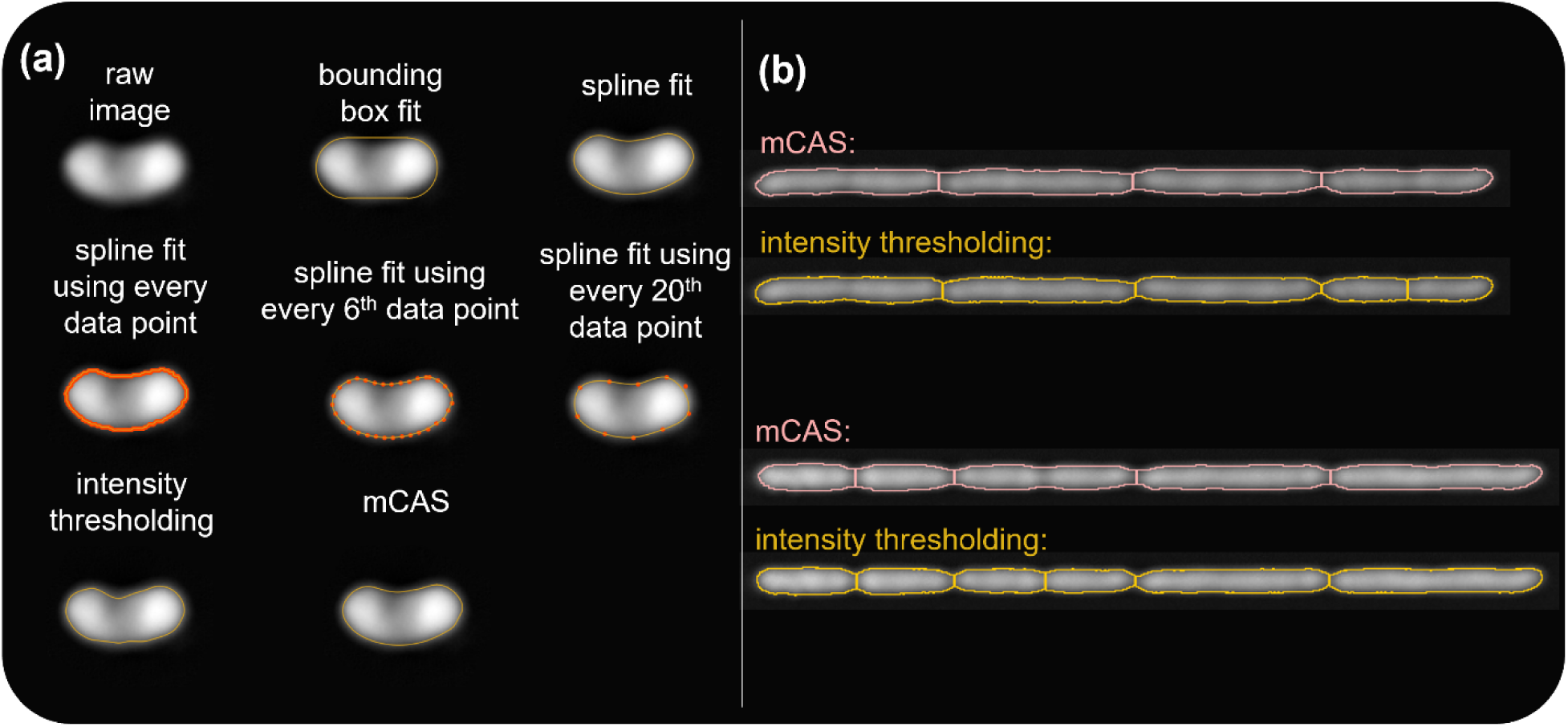
Comparison of segmentation approaches. **(a)** *Top row*: segmentation of *M. extorquens* using bounding box fitting versus spline fitting. *Middle row*: examples of spline fitting at different sampling rates (every data point, every 6^th^ point, every 20^th^ point). *Bottom row:* comparison of intensity thresholding with spline fitting versus mCAS with spline fitting for a single *M. extorquens* cell. **(b)** Examples of *E. coli* segmentation errors from watershed separation applied to intensity-thresholded masks, contrasted with accurate mCAS-derived contours.

### mCas Validation with E. coli

To validate the accuracy of mCAS-generated masks from QPI, we first performed a direct comparison with standard fluorescence-based segmentation. Specifically, we imaged live *E. coli* (63ME121R) expressing the tdCherry fluorescent protein [31] and generated cell masks from the corresponding fluorescence images using optimized intensity-thresholding. When compared, the fluorescence-derived masks and QPI-derived mCAS masks showed strong agreement as highlighted by the high (> 90%) accuracy and Dice scores for n = 350 (**Fig. 2b**). This strong agreement demonstrated that mCAS applied to QPI faithfully reproduced the accepted cell boundaries defined by fluorescence imaging, confirming its reliability as a label-free segmentation approach.

### Application of mCAS to Growing E. coli

We next evaluated the performance of mCAS on time-lapse QPI during *E. coli* growth. Specifically, we analyzed cells growing in optically transparent one-dimensional (1D) microarrays, which we recently showed preserve phase information with 60% higher accuracy compared to two-dimensional (2D) confinement. [23] Using mCAS, we segmented individual cells with two iterations of ImageJ’s built-in binary erosion function, followed by adjustable watershed segmentation at σ = 1.5. For shape fitting, we used the bounding box method, which matches the rod-like morphology of *E. coli*.

We found that mCAS substantially reduced common segmentation errors in comparison to conventional intensity-thresholding combined with watershed. Specifically, we found that thresholding often produced two types of artifacts: **(1)** artificial fragmentation of single cells into multiple regions of interest (ROIs) (**Fig. 3b**, **Fig. S2**), and **(2)** overlapping ROIs after bounding box shaping. Overlaps typically arose from diagonal boundaries introduced during watershed processing of thresholded images, which can incorrectly merge adjacent cells into overlapping bounding boxes. By contrast, mCAS-generated masks rarely exhibited such artifacts, thereby minimizing false overlaps (**Fig. S3**). mCAS also reduced over-segmentation. By providing more coherent and biologically accurate cell boundaries, mCAS produced labeled masks that were directly compatible with downstream lineage-tracking tools such as *Lineage Mapper* [32] (**Fig. 4**). This improved both the fidelity of the segmentation and overall analysis throughput.

**Figure 4:**
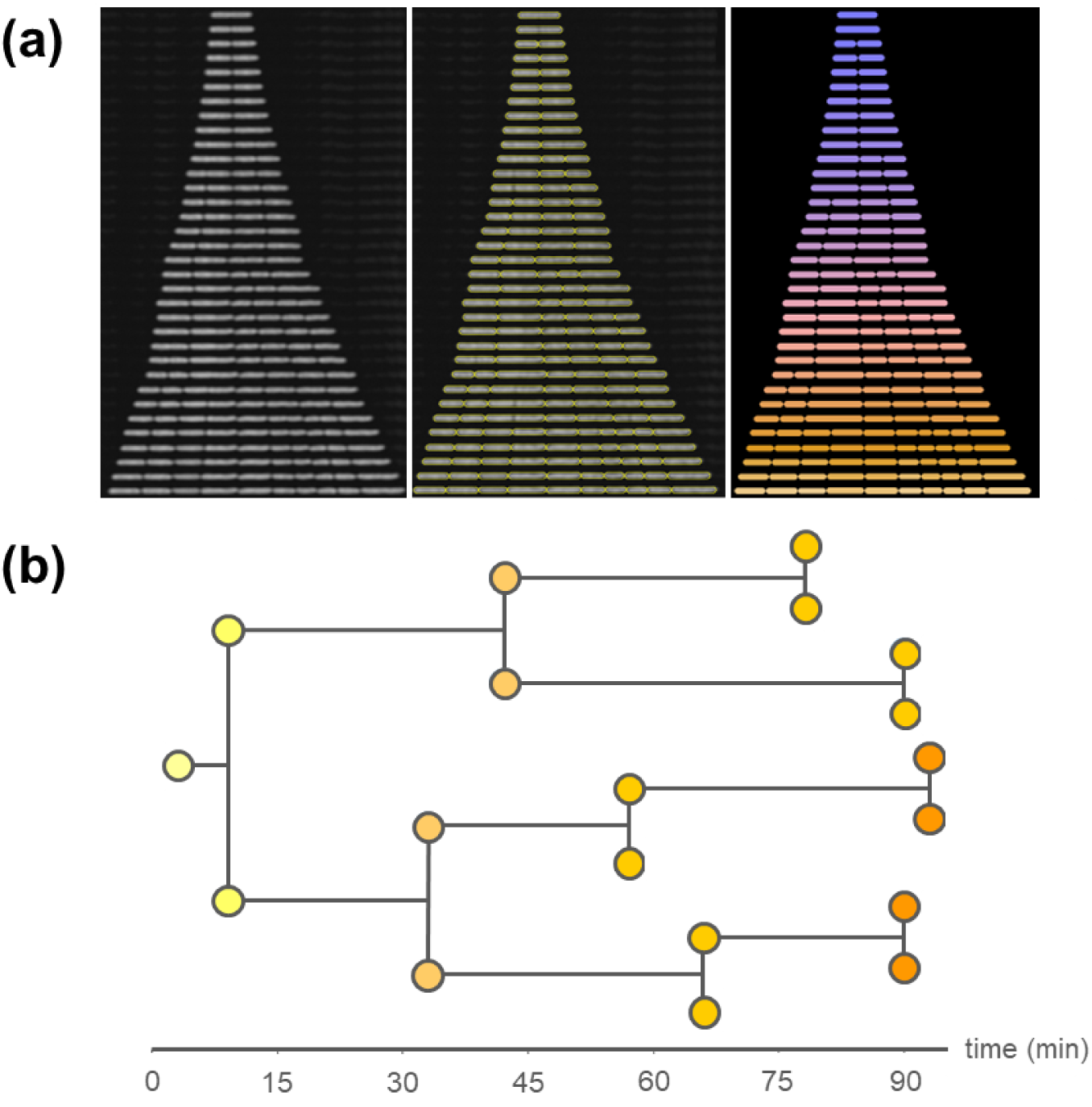
Single-cell tracking using mCAS segmentation. **(a)** *Left:* timelapse montage of *E. coli* growth in a 1D microfluidic device imaged by QPI. *Middle:* mCAS-derived regions of interest (ROIs) overlaid on the same montage. *Right:* labeled masks generated from mCAS segmentation. **(b)** Lineage map constructed from the labeled mCAS masks, showing cell divisions over time.

### Segmentation of M. extorquens

Unlike the simple rod morphology of *E. coli*, some cell types contain complex intracellular structures that complicate segmentation. In such cases, traditional intensity-based methods often fail to delineate cell boundaries due to heterogeneous internal contrast. To test mCAS under these conditions, we analyzed *M. extorquens*, which accumulates polyhydroxybutyrate (PHB) granules under nutrient limitation. [33] The presence of PHB produced strong variations in optical density within cells, making accurate segmentation particularly challenging.

By combining edge detection, watershed segmentation, and spline-based contour shaping, mCAS overcame these challenges and produced masks with high boundary fidelity. We benchmarked mCAS against manually segmented reference masks from 100 *M. extorquens* cells. The choice of n = 100 cells was based on error convergence analysis, in which mean Dice and accuracy scores for mCAS-segmented cells remained stable for varying number of cells (**Fig. 5**). mCAS consistently outperformed Otsu thresholding with spline fitting, achieving average Dice and accuracy metrics exceeding 0.98 (**Fig. 5**). The resulting precision enabled reliable delineation of cells with complex intracellular features, supporting accurate downstream quantitative analysis.

**Figure 5:**
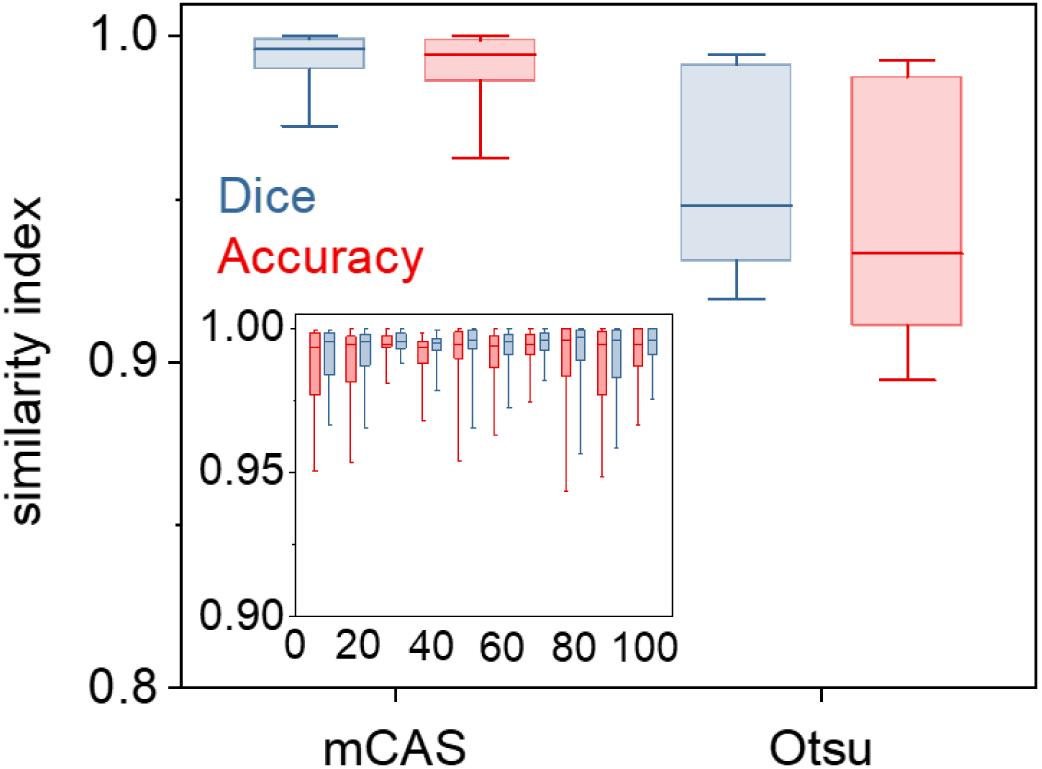
mCAS performance. Box charts of the Dice and accuracy metrics comparing n = 100 masks generated by mCAS and Otsu’s thresholding for *M. extorquens*. *Inset* shows Dice and accuracy scores for mCAS segmentation over increasing sample sizes. Box charts represent 25-75% of the individual scores; whiskers indicate 10-90% of individual scores; horizontal lines indicate the median score

### Comparison of mCAS with Deep Learning Methods

DL has transformed biological and biomedical imaging, driving major advances in image processing, cell segmentation, and clinical diagnostics. Well-trained DL models can achieve near-human precision, often reducing the need for manual curation. [8,34–37] DL has also been applied extensively to QPI, enabling label-free segmentation and classification of diverse cell types and organelles. [14, 38] Despite these strengths, AI segmentation has important limitations. Non-foundational networks, such as conventional convolutional neural networks (CNNs), often lack generalizability and perform best on datasets closely resembling their training data. Adapting such models to new imaging modalities or different cell types typically requires re-training and fine-tuning. [38] Foundational networks such as Meta’s *Segment Anything Model* (SAM) [39] offer broader applicability, but they still rely on user input for refinement and often require post-processing to improve boundary accuracy. While operations such as binary erosion can mitigate boundary artifacts, they frequently distort shapes or shrink masks, reducing agreement with ground truth annotations.

In contrast, mCAS requires no training data, avoiding the significant time and computational cost associated with DL model development. For direct comparison, we trained our previously reported U-Net model [38] using 400 mCAS-generated 256×256 images of single *M. extorquens* on a CPU, which required ∼40 hours. Training-set size was optimized through experiments with smaller datasets and augmentation (**Fig. S4**). While the trained U-Net outperformed Otsu thresholding, it did not achieve the segmentation quality of mCAS when benchmarked against 100 manually segmented images (**Fig. 6a**). Moreover, the U-Net exhibited slower processing times than mCAS implemented in ImageJ (**Fig. 6b**) and struggled with delineating adjacent or overlapping cells (**Fig. 6c**).

**Figure 6:**
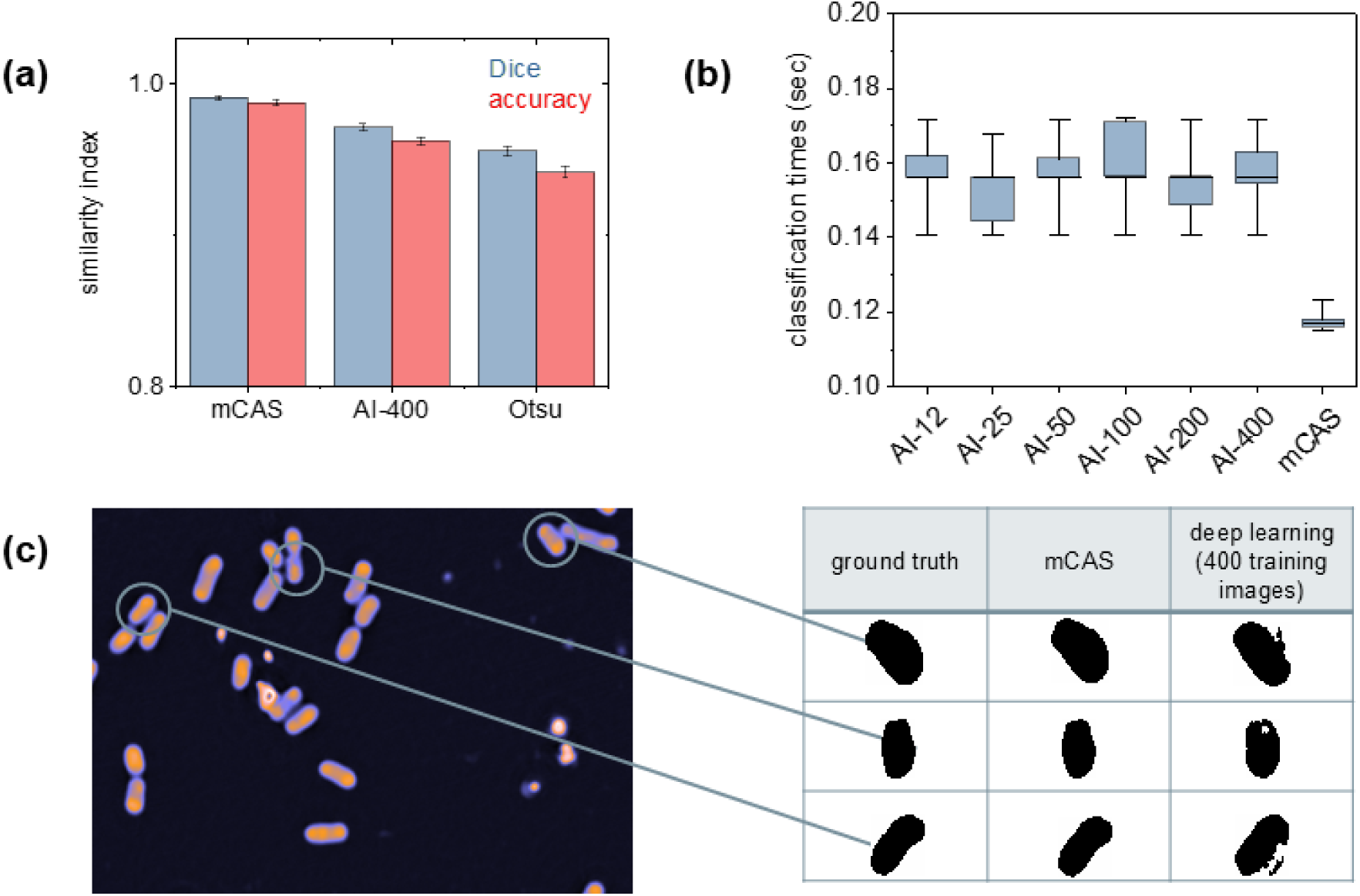
Performance comparison between mCAS and AI-based segmentation of *M. extorquens*. **(a) Mean Dice and accuracy scores for 100 segmentation masks from each method compared with corresponding manually segmented masks. Error bars show standard error of the mean. (b) Classification times for 100 test images using the mCAS method versus a U-Net model trained on 12, 25, 50, 100, 200, or 400 images. Box plots show the 27-75% range, whiskers show the 10-90% range, and median scores are indicated by a horizontal line. (c) Examples of poor DL segmentation compared with CAS and ground truth.**

We also evaluated SAM for QPI segmentation of fixed *M. extorquens* using the online demonstration software. This approach required manual input for each cell and frequently produced inaccurate masks for both *E. coli* and *M. extorquens* (**Fig. S5**). Together, these results demonstrate that mCAS provides a computationally efficient, cost-effective, and highly reliable alternative to both foundational and non-foundational DL methods, particularly in settings where training data are limited or cell morphologies are highly variable.

## Discussion

mCAS addresses a central challenge in QPI: the absence of a generalized, efficient, and accurate segmentation method specifically for microbial cells. Benchmarking against standard intensity-based approaches and state-of-the-art DL models demonstrate that mCAS consistently achieves higher accuracy, faster processing, and broader adaptability across diverse microbial morphologies. In contrast, most existing QPI-based segmentation approaches were developed for mammalian cells. For example, a representative 2D method achieved computation times of approximately 1 second per cell, [19] while CellSNAP, a 3D approach implemented through CellProfiler, reached similar accuracy at about 2 seconds per cell. [18] By comparison, mCAS is purpose-built for microbial cells, approximately 10-100× smaller than mammalian cells [40], and segments them in just 0.12 seconds per cell. Because microbes are near the resolution limit of most imaging systems, mCAS is specifically optimized for this challenging regime.

Unlike DL pipelines that require large, curated datasets and significant computational resources, mCAS generalizes effectively without retraining [8,34–37]. Even well-trained networks often perform optimally only on datasets that closely resemble their training data [38], and foundational models such as SAM [39] still depend on manual input and post-processing. In contrast, mCAS achieves Dice and accuracy scores above 0.98 directly from raw QPI data, eliminating these bottlenecks while enabling high-throughput imaging. Biological validation further underscores this versatility. In *E. coli*, mCAS reduced over-segmentation and overlapping ROI artifacts commonly observed in thresholding-based pipelines, producing masks directly compatible with lineage-tracking tools. In *M. extorquens*, where intracellular PHB granules create heterogeneous optical contrast, mCAS outperformed Otsu thresholding with spline fitting, delivering reliable contours even under challenging conditions. Together, these results demonstrate that mCAS maintains robust performance across diverse microbial morphologies and intracellular complexities.

Computational efficiency represents another major strength of mCAS. The method can segment more than 1,000 single-cell QPI images in under two minutes using standard desktop hardware. This rapid processing eliminates a major bottleneck in QPI workflows, enabling large-scale downstream analyses such as lineage mapping, growth tracking, and dry-mass quantification without the delays associated with DL training or manual correction. Overall, mCAS provides a training-free, computationally efficient, and biologically adaptable solution for microbial segmentation in QPI. By combining edge detection with modular post-processing, it bridges a key gap between raw label-free imaging and quantitative single-cell biology. Looking forward, extending mCAS to higher-order organisms and multidimensional or integrative imaging modalities [41] will establish it as a foundation for real-time QPI-based phenotyping and further broaden its impact in quantitative cell biology.

## Acknowledgments

We gratefully acknowledge funding from the US Department of Energy, Office of Science, Office of Biological and Environmental Research (DE-SC0022318 and DE-SC0022282) for supporting this work. We also acknowledge fruitful discussions with Dev Shrestha.

## Contributions

SJT performed all image processing described in this work, as well as the *M. extorquens* and 63ME121R experiments; SN performed all DH5α experiments; ABA and CJM provided biological materials; AEV supervised the research and wrote the manuscript with SJT.

## Data Availability Statement

Data are available from the corresponding author upon request, while all algorithms developed for this work are available as Supplementary Information.

## Methods Strains

To verify the generality of our results, we imaged two types of bacteria, namely: *Escherichia coli* DH5α and 63ME121R [31], and *Methylobacterium extorquens* PA1 [42] (CM2730 [43]).

## Growth/Fixation Conditions

### Escherichia coli

All *E. coli* strains were grown in a water-bath incubator (C76, New Brunswick Scientific) at 37 °C, 180 rpm, using Mueller-Hinton broth (Difco 275730, BD). The DH5α and 63ME121R strains were transferred from glycerol stocks at -80 °C, inoculated into 5 mL of medium (round-bottom polystyrene tubes, VWR) and grown to early stationary phase. Cultures were then diluted to an optical density (at λ = 600 nm) of 0.01 in 20 mL fresh medium (125 mL glass flasks, Corning) and incubated overnight (12 h, 37 °C, 180 rpm). The following day, they were again diluted to an OD of 0.01 in 20 mL medium and grown to mid-exponential phase. Imaging was performed on samples from this mid-exponential phase.

### Methylobacterium extorquens

*M. extorquens* was cultured in MPIPES medium [43] with 7 mM succinate at 28 °C in a shaker incubator (Excella E24, New Brunswick Scientific). Cells were transferred from agar plates (stored at 4 °C) into 5 mL of medium and grown for 24 h before inoculation into 20 mL fresh medium (125 mL glass flasks, Corning). For fixed-cell imaging, cultures were harvested at 17, 21, and 48 h, fixed in 2% paraformaldehyde (2 h), washed three times in PBS, and stored at 4 °C.

### 1D assays

Single *E. coli* cells were confined in one-dimensional (1D) microarrays, as described previously [23], covered by a nutrient-doped membrane. The microarrays were fabricated by electron-beam lithography in SU8, transferred to PDMS, and subsequently into a UV-curable polymer index-matched to water (Bio-133, My Polymers). A polymer film thickness of ∼500 µm after imprinting was achieved by depositing ∼100 µL of liquid prepolymer on the PDMS stamp, enabling imaging with a 63× objective (NA 0.7, PH2). Nutrients were delivered through the integrated agarose membrane following established protocols [23,44].

#### Imaging *E. coli* in 1D and 2D

For 1D confinement, we acquired 3D QPI stacks with a 63× objective (NA 0.7, PH2; vertical step size 300 nm) using a CCD camera (GS3-U3-28S4M, Point Grey Research; 3.65 µm pixel size), including halo effect correction [22–23]. Multiple fields were imaged every ∼3 min (Metamorph, Molecular Devices). For 2D confinement, *E. coli* cells were deposited on nutrient-containing agarose pads (1.5% UltraPure agarose, Invitrogen, in Mueller-Hinton broth). Pads were prepared by sandwiching ∼300 µL of molten agarose-broth mixture between two coverslips (25 × 50 mm). After a 20 min drying period at room temperature (∼90 µm thickness), cells were deposited, air-dried for 5 min, and covered with a coverslip. Fluorescence images were acquired under incoherent excitation centered at 560 nm.

#### M. extorquens imaging

Fixed *M. extorquens* cells were imaged in 3D using a 100× oil immersion objective (NA 1.3, PH3; Leica) with a 150 nm vertical step size and a CCD camera (Hamamatsu Orca Flash 4.0; 6.5 µm pixel size). Cells were mounted on 2% agarose pads (∼90 µm thick) prepared as above.

### Code

We used the UNET code created by Sheneman et al., which is available at https://github.com/sheneman/deep_lipid. [38]

### Training Set

Training images (256 × 256 pixels) were generated from CAS-segmented *M. extorquens* images. Sets of 12, 25, 50, 100, 200, and 400 unique images were sampled from a pool of 2,400.

## Data Augmentation

For augmented models, training images were doubled using random rotation, flipping, and scaling (± 30%).

## Metrics

Segmentation performance was evaluated using the Dice–Sørensen coefficient and accuracy metric:

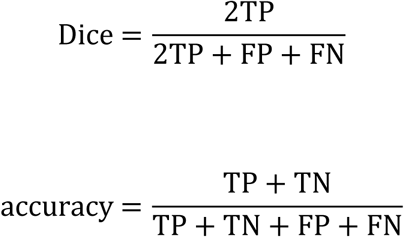

where TP and TN denote true positives and true negatives, and FP and FN denote false positives and false negatives, respectively.

